# Sleep-like bistability, loss of causality and complexity in the brain of Unresponsive Wakefulness Syndrome patients

**DOI:** 10.1101/242644

**Authors:** M. Rosanova, M. Fecchio, S. Casarotto, S. Sarasso, A.G. Casali, A. Pigorini, A. Comanducci, F. Seregni, G. Devalle, O Bodart, M. Boly, O. Gosseries, S. Laureys, M. Massimini

## Abstract

Unresponsiveness Wakefulness Syndrome (UWS) patients may retain intact portions of the thalamocortical system that are spontaneously active and responsive to sensory stimuli. In these patients, Transcranial Magnetic Stimulation combined with electroencephalography (TMS/EEG) also reveals preserved cortical reactivity, but in most cases, the residual thalamocortical circuits fail to engage complex causal interactions, as assessed by the perturbational complexity index (PCI).

Another condition during which thalamocortical circuits are intact, active and reactive, yet unable to generate complex responses, is physiological non-rapid eye movement (NREM) sleep. The underlying mechanism is bistability: the tendency of cortical neurons to fall into a silent period (OFF-period) upon receiving an input.

Here we tested whether a pathological form of bistability may be responsible for loss of brain complexity in UWS patients. Time-frequency decomposition analysis of TMS/EEG responses in UWS patients revealed the occurrence of OFF-periods (detected as a transient suppression of high-frequency oscillations in the EEG) similar to the ones evoked by TMS in the cortex of healthy sleeping subjects. Pathological OFF-periods were detected in any cortical area, significantly impaired local causal interactions (as measured by PLF) and prevented the buildup of global complexity (as measured by PCI) in the brain of UWS patients.

Our results draw a first link between neuronal events (OFF-periods) and global brain dynamics (complexity) in UWS patients. To the extent that sleep-like bistability represents the common functional endpoint of loss of complexity, detecting its presence and tracking its evolution over time, may offer a valuable read-out to devise, guide and titrate therapeutic strategies aimed at restoring consciousness.

## Introduction

Patients diagnosed with unresponsive wakefulness syndrome (UWS), previously known as Vegetative State *(1)*, recover sleep-wake cycles, can open their eyes, but do not show behavioral signs of consciousness *(2)*. Despite behavioral unresponsiveness, many of these patients retain large parts of the thalamo-cortical system that are structurally intact, spontaneously active *(3–5)* as well as reactive to sensory stimuli, though cortical responses tend not to propagate beyond primary areas *(3, 6–9)*. Preserved cortical reactivity in UWS can be directly demonstrated by Transcranial Magnetic Stimulation in combination with the electroencephalogram (TMS/EEG); apart from severe post-anoxic patients, TMS always elicits significant cortical responses in UWS patients. In a minority of such patients, EEG responses to TMS are similar to those observed in healthy awake subjects, suggesting that they may retain a covert capacity for consciousness. However, in most cases, the EEG response to TMS is simple and stereotypical: specifically, direct cortical perturbation elicits a strong initial activation, which fails to engage in sustained, complex patterns of interactions, as assessed by the perturbational complexity index (PCI) *(10)*. Thus, in many UWS patients cortical circuits seem to be active, reactive but blocked in a pathological low-complexity state.

NREM sleep is a physiological condition in which thalamocortical circuits are structurally intact, functionally active and reactive, yet unable to engage in long-range, complex responses *(11, 12)*. A recent study employing intracortical single pulse electrical stimulation (SPES) and simultaneous local field potential (LFP) recordings in humans *(13)*, suggests that the mechanism responsible for this impairment in NREM sleep is attributed to cortical bistability, which is the tendency of cortical neurons to fall into an OFF-period in response to transient increases in activity *(14, 15)*. During physiological sleep, in silico, in vitro as well as in vivo models suggested activity-dependent K+ currents *(16, 17)* as well as active inhibition *(18, 19)* as putative determinants for cortical bistability. SPES and LFP recordings show that, due to bistability, neurons react briefly to incoming signals and then fall into an OFF-period, which rapidly disrupts the effects of the initial input. Thus, in physiological sleep this simple dynamics leads to a breakdown of causal interactions among cortical areas and prevents the emergence of sustained, complex patterns of interaction, despite preserved activity and reactivity.

Can a pathological form of bistability also play a role in the residual cortex of low-complexity UWS patients? Asking this question is relevant for at least two reasons. First, bistability represents a basic default mode of cortical activity *(20)*, which can be engendered by physiological changes as well as by pathological alterations, such as shifts of the inhibition/excitation balance *(21)* or white matter lesions *(22)*. Second, neuronal bistability can disrupt complex cortico-cortical interactions, but is in principle reversible.

Here, we specifically test the following hypotheses: (i) pathological sleep-like bistable dynamics occur in the cortex of awake UWS patients and (ii) these dynamics are responsible for the collapse of causality and overall brain complexity associated with loss of consciousness following brain injury. To do so, we analyzed TMS-evoked EEG potentials (TEPs) recorded in low-complexity UWS patients with the same analysis previously used on intracranial SPES-evoked LFP potentials during sleep *(13)*. First, we show that in UWS patients with their eyes open, the EEG response to TMS in anatomically preserved cortical areas matches the electrophysiological criteria for bistability, as assessed during NREM sleep, i. e. a simple positive-negative wave, associated with a neuronal OFF-period. Then, we demonstrate that pathological, sleep-like bistable dynamics rapidly disrupt local causal effects of TMS (as indexed by phase-locking measures) and in turn, the emergence of global complex cortico-cortical interactions (as indexed by PCI) in the brain of UWS patients.

## Results

We analyzed 72 TMS/EEG measurements performed in 16 awake UWS patients and 20 healthy subjects during wakefulness and NREM sleep, while stimulating both frontal and parietal cortex. Specifically, we assessed (1) the presence of cortical OFF-periods, i.e. the occurrence of a TMS-evoked slow wave (< 4 Hz) associated with a significant high frequency (> 20 Hz) suppression of EEG power compared to pre-stimulus, (2) the impact of the OFF-periods on local causal interactions by means of broadband (> 8 Hz) phase-locking factor (PLF), (3) the consequences of the OFF-period on the build-up of complex global interactions as indexed by the time course of PCI. For a detailed description of the experimental and analytical procedures, see the Materials and Methods section and Fig. S1.

### TMS reveals sleep-like OFF-periods in the cortex of awake UWS patients

TEPs recorded in UWS patients consisted of a slow wave, which was associated with an initial activation rapidly followed by a significant high frequency (> 20 Hz) suppression of EEG power (HFp) starting at around 103±9 ms (mean±SEM; Fig. 1B and Fig. S2B). This pattern of local reactivity matches the criteria for an OFF-period *(13, 23–26)* and was observed in all stimulated areas (both frontal and parietal bilaterally; Fig. 2A) in each of the 16 UWS patients, irrespective of the presence/absence of spontaneous slow waves in the ongoing pre-stimulus activity (Fig. 2B). The responses found in UWS patients differed markedly from awake healthy subjects stimulated over the same areas (Fig. 1A and Fig. S2A); in this latter case, evoked slow waves were absent, low-frequency (< 4 Hz) EEG amplitude (max SWa, see Materials and Methods) was significantly lower (Bonferroni corrected p<0.05 for parietal stimulation, and p<0.01 for frontal stimulation; Fig. 1D and Fig. S2C, top panel) and the suppression of high-frequency power was never observed (Fig. 1D and Fig. S2C, middle panel). Conversely, the UWS response was similar to the one found in healthy subjects during NREM (Fig. 1C), where TMS evoked a slow wave with a comparable level of low-frequency EEG amplitude (Bonferroni corrected p=0.36; Fig. 1D, top panel) associated with a significant high-frequency suppression (Bonferroni corrected p=0.99 Fig. 1D, middle panel) starting at around 127±11 ms.

**Fig. 1.**
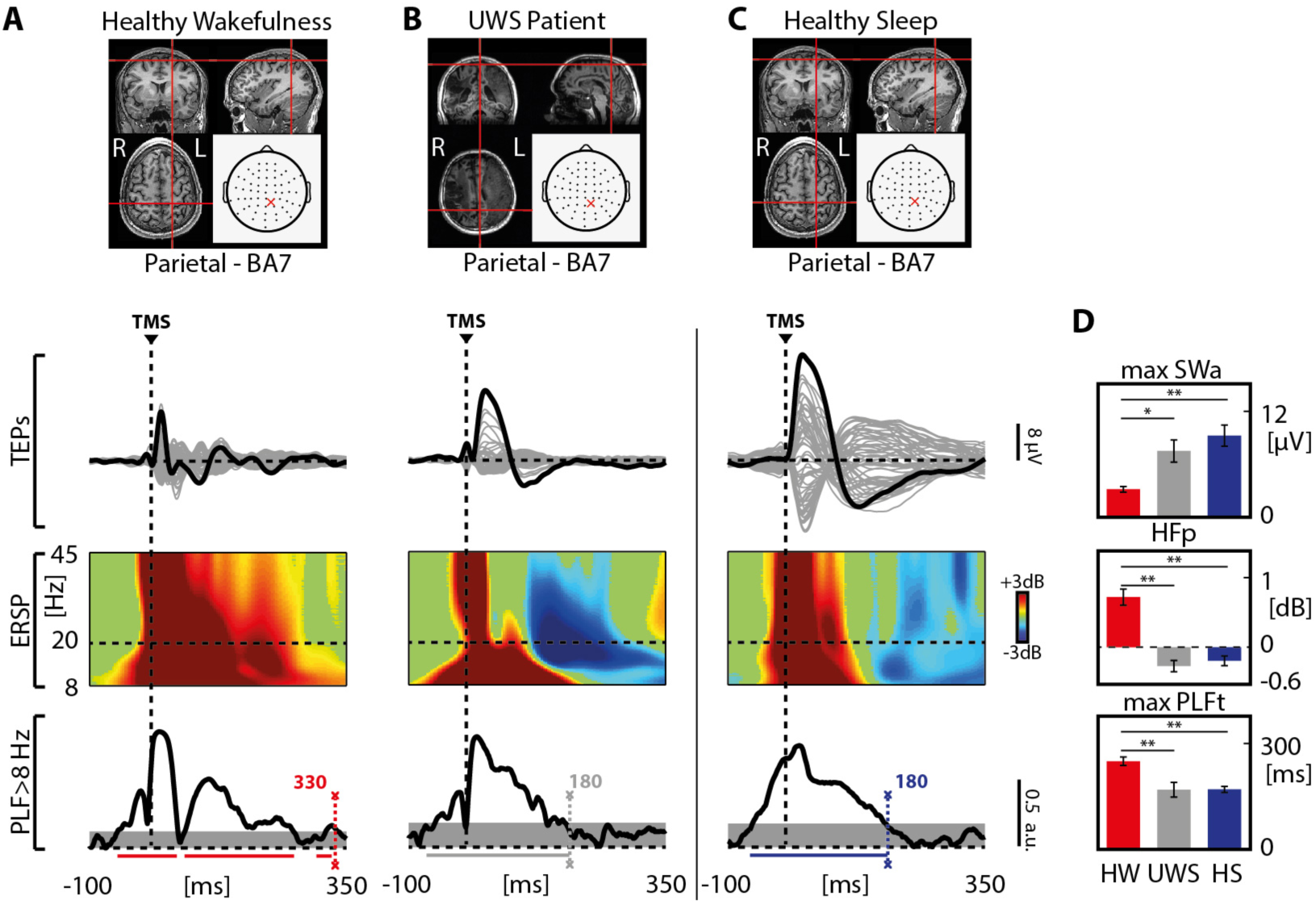
TMS evokes a stereotypical sleep-like response that is associated with high-frequency (> 20 Hz) suppression and with an early drop of PLF in UWS patients. Results for a representative healthy subject stimulated during wakefulness (HW) and during NREM sleep (HS) and a representative Unresponsive Wakefulness Syndrome patient (UWS) are shown. **(A to C)** Structural MRIs (T1-weighted) and TMS cortical targets as estimated by NBS are shown (top). TMS-evoked EEG responses recorded at all 60 channels (grey traces) are also depicted in the butterfly plots. The thicker black trace highlights the electrode with the larger TEP. For this electrode the ERSP and the PLF calculated above 8 Hz are presented. For the ERSP, significance for bootstrap statistics is set at α < 0.05 (absence of any significant activations is colored in green): statistically significant increases of power compared to baseline are colored in red, while blue represents significant power decreases. The dashed horizontal line indicates the 20 Hz frequency bin. For the PLF, statistical differences (α < 0.01) compared to the baseline (from -500 to -100 ms) were assessed and the timing of the last significant time point was considered (indicated by the light-dashed vertical line). Time points above statistical threshold (grey shaded area) are underlined by a horizontal line (red, gray, blue for HW, UWS, HS, respectively). A dashed vertical line marks the occurrence of the TMS pulse. **(D)** Group-level comparison among awake healthy subjects (red), awake UWS patients (gray) and healthy subjects during NREM sleep (blue) of the slow wave amplitude (max SWa, Kruskal-Wallis test, p = 0.7*10^−3^), of the high frequency (> 20 Hz) power averaged between 100 and 350 ms (HFp, Kruskal–Wallis test, p = 0.2*10^−6^) and of the duration of PLF (max PLFt, Kruskal-Wallis test, p = 0.4*10^−3^) are shown. In each histogram, the group mean and standard error are reported for each measure, together with the statistical significance of post-hoc comparisons between groups (Wilcoxon ranksum test, * p < 0.05, ** p < 0.01).

**Fig. 2.**
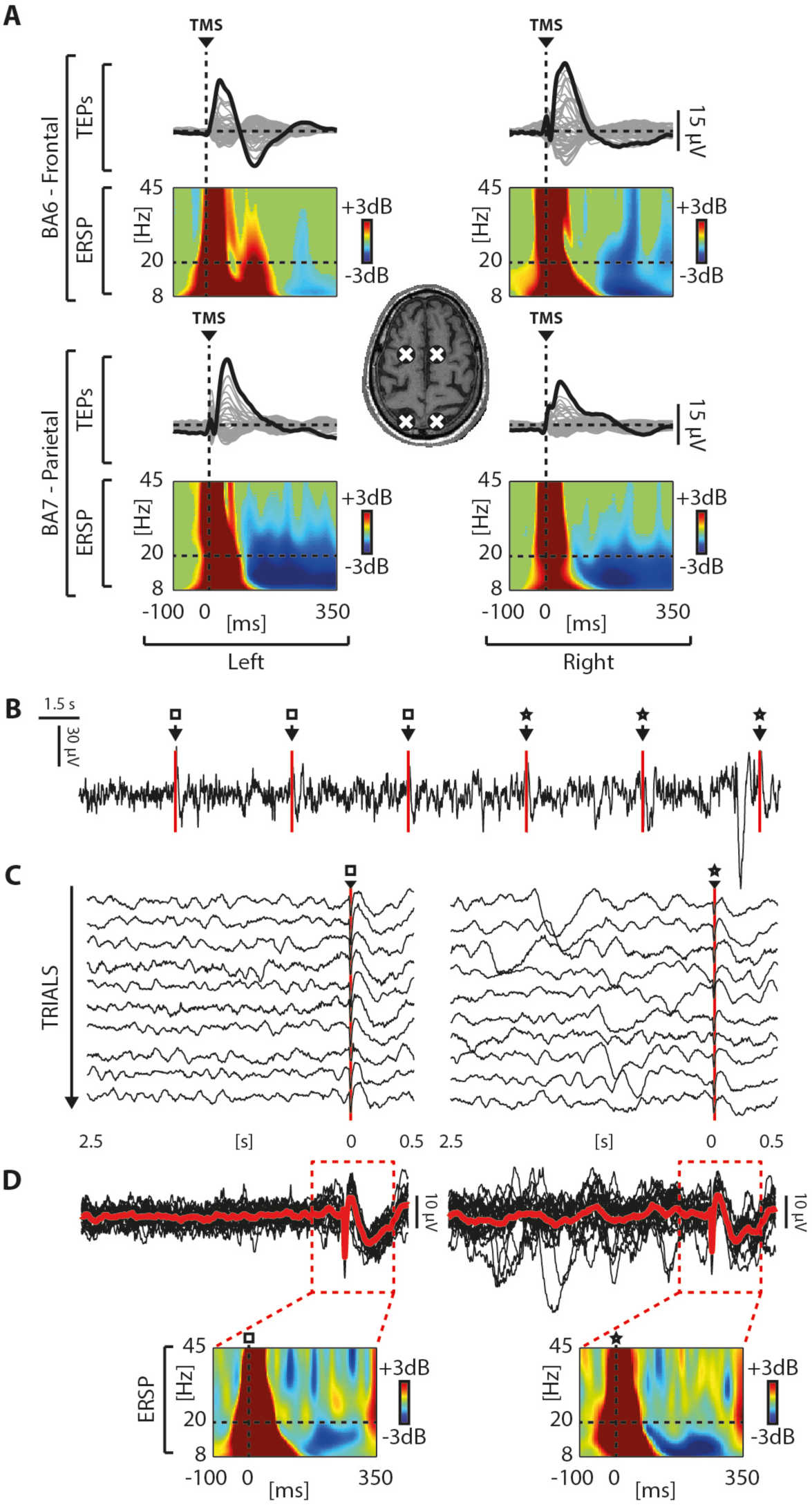
TMS evokes a stereotypical sleep-like response and a high-frequency (> 20 Hz) suppression irrespective of stimulation site and pre-stimulus activity. **(A)** White crosses on the structural MRI (T1-weighted) indicate the cortical TMS targets (Frontal/Parietal and Left/Right) in patient 15. For each cortical target TMS-evoked potentials recorded from all 60 channels (grey traces) are shown and the electrode with the larger TEP is highlighted (black trace) with the corresponding ERSP. Significance for bootstrap statistics in the time-frequency power spectra is set at α < 0.05: power increase compared to the baseline is colored in red, power decrease in blue and the non-significant values are colored in green. The dashed horizontal line marks the 20 Hz frequency bin and the dashed vertical line indicates the occurrence of the TMS pulse. **(B)** EEG activity (one representative electrode re-referenced to the mathematically-linked mastoids - Cz) recorded in patient 4 while TMS was delivered with an inter-stimulus interval randomly jittering between 5000 and 5300 ms. Empty squares and stars indicate TMS pulses delivered over an ongoing activity not showing (empty squares) or showing (stars) spontaneous slow waves. **(C)** Similar to (B), the empty square and the star indicate trials in which TMS pulses were delivered over an ongoing activity respectively not showing or showing spontaneous slow waves. **(D)** The same trials shown in (C) are overlapped, averaged (red lines) for both conditions and the corresponding ERSPs are shown in the bottom panels. Color coding is the same of (A).

### OFF-periods disrupt local causality in the cortex of awake UWS patients

The duration of the causal effects of TMS on local cortical activity, as assessed by the PLF, was short-lived in UWS patients. Indeed, the latest significant PLF value (max PLFt, see Materials and methods) occurred at 167±21 ms when stimulating parietal cortex and at 188±18 ms when stimulating frontal cortex (Fig. 1D, Fig. S2C). These values roughly corresponded to the timing of the maximum of high frequency (> 20 Hz) suppression (max SHFt) and were similar to the max PLFt of healthy controls during NREM sleep (168±9 ms - Fig. 1D). On the contrary, in healthy awake controls, PLF persisted until 248±12 ms when stimulating parietal cortex and 248±15 ms when stimulating frontal cortex (Fig. 1D, in Fig. S2A). These results were statistically significant at the group level, whereby max PLFt was significantly shorter in UWS patients and healthy subjects during NREM sleep in comparison to healthy awake subjects (Bonferroni corrected p<0.01 for parietal stimulation, and p<0.01 for frontal stimulation; Fig. 1D and Fig. S2C, bottom panel).

Next, we asked whether the three distinctive features of the cortical response found in UWS (i.e. the presence of a slow wave-like response, high frequency (> 20 Hz) suppression and shorter PLF duration) were related. These variables are thought to reflect neurophysiological events (such as the level of neuron membrane polarization, the level of neuronal silencing and its impact on deterministic responses) that are causally linked and showed significant correlation in previous intracranial *(13)* and extracranial *(27)* studies. In order to demonstrate this relationship, we computed linear correlations between max SWa and the maximum level of high frequency (> 20 Hz) suppression (max SHFp) and between the timing of the maximum SHFp (max SHFt) and max PLFt, respectively. Interestingly, max SWa was significantly correlated with max SHF (R^2^= 0.4, p < 0.01; Fig. 3, left). Also, max SHFt was significantly correlated with max PLFt (R^2^ = 0.34, p < 0.01; Fig. 3, right), showing that (i) larger evoked slow waves corresponded to more pronounced OFF-periods and (ii) earlier OFF-periods corresponded to an earlier dampening of the causal effects induced by the perturbation.

**Fig. 3.**
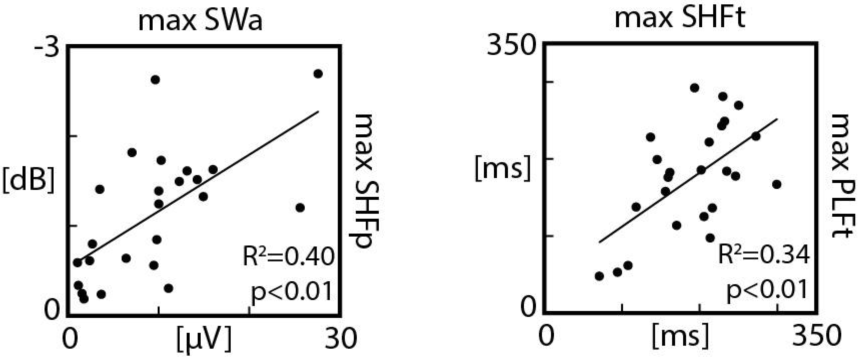
Slow-wave amplitude, high frequency (> 20 Hz) suppression of power and phase-locking factor are correlated in UWS patients. On the left, the correlation between the maximal amplitude of the evoked slow wave (max SWa) and the maximal level of high-frequency power suppression (max SHFp) is shown. On the right, the correlation between the timing of the maximum high frequency (> 20 Hz) suppression (max SHFt) and the latency at which PLF fell below the threshold for significance (max PLFt) is shown. For both correlations, the coefficient of determination R^2^ and the significance level p are reported.

### OFF-periods reduce global complexity

We finally asked whether cortical OFF-periods and their aftermath on local causality might be responsible for the loss of global brain complexity. All UWS patients included in the present study were characterized by levels of brain complexity (PCI range: 0.13-0.30) invariably lower than the ones measured in healthy awake subjects (PCI range: 0.32-0.64). These lower PCI values could be explained by a difference in the time-course of the build-up of brain complexity after TMS (PCI(t), see Materials and methods). While in awake healthy subjects PCI(t) kept growing up to about 300 ms (272±4.6 ms, Fig. 4A, right plot), in low complexity UWS patients PCI(t) grew initially but reached a plateau at an earlier time point (197± 12 ms) resulting in a significantly shorter build-up (p<0.01). Crucially, the timing at which global complexity stopped growing (max PCIt) showed a significant positive correlation with the timing of the OFF-period (max SHFt; R^2^ = 0.46, p < 0.01; Fig. 4C, upper plot) as well as with the timing at which local causality broke-off (max PLFt; R^2^ = 0.56, p < 0.01; Fig. 4C, lower plot). This result is highlighted in Fig. 4B for a representative UWS patient (patient n. 12), where the time courses of high-frequency EEG power modulation, broadband PLF and PCI are depicted. To further strengthen the link between OFF-periods, loss of local causality and global complexity, we observed that recovery of consciousness (as assessed by the CRS-R) in a longitudinally recorded patient (Patient 16) was paralleled by a progressive reduction of high-frequency (>20 Hz) suppression, a concurrent prolongation of PLF and an increase of PCI up to values found in conscious awake subjects. (Fig. 5).

**Fig. 4.**
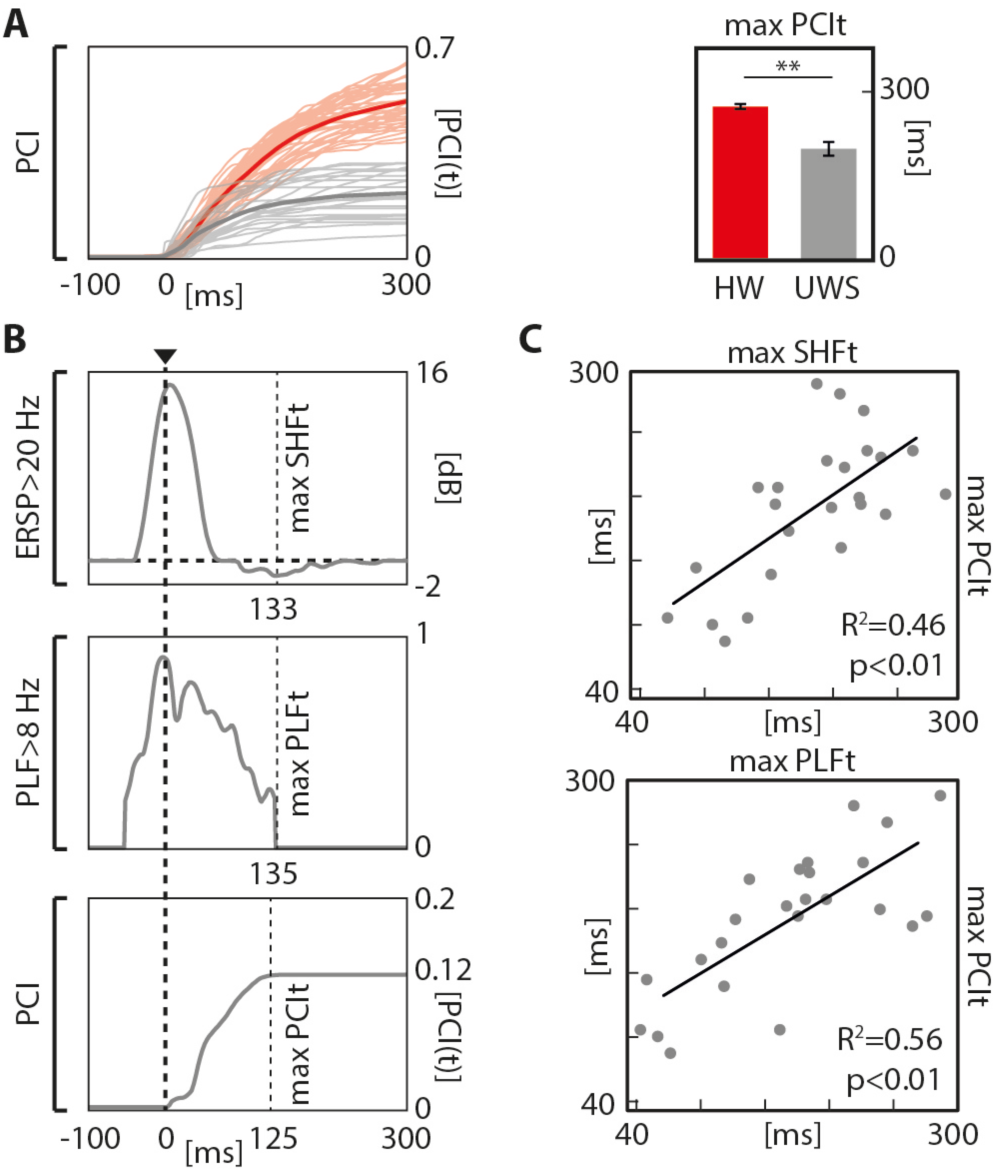
The occurrence of an OFF-period can prevent the build-up of PCI. **(A)** For each TMS/EEG measurements, the temporal evolution of PCI, PCI(t), calculated in awake healthy subjects (light-red lines) and UWS patients (light-gray lines) are shown together with their grand average (thick red and thick gray lines, respectively). On the right, histograms indicate the grand average of the time at which PCI reach its maximum value (max PCIt), together with the standard error and the statistically significant differences (Wilcoxon ranksum test, ** p < 0.01). **(B)** Top and middle panels show, for one channel close to the stimulation site in one representative UWS patient, the time course of the averaged high frequency (>20 Hz) power and the time course of the significant PLF above 8 Hz, respectively. Bottom panel shows, for the same representative UWS patient, the temporal evolution of PCI. The light-dashed vertical lines mark the exact timing of the maximum high frequency (> 20 Hz) suppression (max SHFt, top), the latency at which PLF fell below the threshold for significance (max PLFt, middle) and the time at which PCI reach its maximum value (max PCIt, bottom), respectively. The bold dashed vertical line indicates the onset of the TMS pulse. **(C)** The correlation between max SHFt and max PCIt (top chart) and the correlation between max PLFt and max PCIt (bottom chart) are shown. For both correlations, the coefficient of determination R^2^ and the significance p level are reported.

## Discussion

Previous studies employing TMS/EEG have shown that most UWS patients retain portions of the cerebral cortex that are active and reactive *(28–31)* but blocked in a state of low complexity *(10, 12)*. Here we investigated the electrophysiological mechanisms underlying this condition; we show that in these low-complexity patients the cortical response to TMS is underpinned by a neuronal OFF-period (the hallmark of cortical bistability). Further, we demonstrate that the occurrence of this OFF-period rapidly disrupts the build-up of causal effects of the initial activation thus preventing the emergence of large-scale complex interactions. Similar electrophysiological events were detected in sleeping healthy controls but were never found in healthy awake subjects, suggesting that a pathological form of bistability may occur in UWS patients. Therefore, the present findings link cortical bistability - a well-characterized neuronal mechanism that is known to play a role in physiological NREM sleep - to the pathophysiology of the UWS.

## Pathological mechanisms of bistability

At the neuronal level, the key feature of bistability in physiological conditions is the occurrence of OFF-periods, reflecting a profound hyperpolarization (down-state) in the membrane of cortical neurons. During NREM sleep, the occurrence of down-states seems to be mainly caused by the enhancement of activity-dependent and leak K+ currents, brought about by decreased levels of neuromodulation from brainstem activating systems *(32–36)* and/or by increased inhibition *(18,19)*. Due to these local dynamics, cortical neurons tend to plunge into a silent, hyperpolarized state, lasting few hundreds of milliseconds, after an initial activation (up-states) *(37, 38)*. In the sleeping brain, the occurrence of synchronous intracellular down-states in cortical neurons is reflected at the extracellular level in large slow waves associated with transient suppressions of high-frequency (>20 Hz) activity that are detectable both at the LFP *(23–26)* and at the EEG level *(27)*. Due to its activity-dependent nature, bistability and the associated down-states can be promptly revealed using a perturbational approach, whereby the impulse-response properties of cortical neurons is probed by means of direct activations. Hence, intracortical stimulations have been employed to investigate bistable dynamics in humans and their effect on the propagation of cortico-cortical evoked potentials during wakefulness and sleep *(13, 39)*.

A key finding of the present work is the demonstration of sleep-like bistable responses in the cortex of UWS patients (Fig. 1B and S2B). Specifically, targeting neuronavigated TMS to intact portions of both their frontal and parietal cortices invariably elicited a stereotypical slow wave associated with a high frequency (> 20 Hz) suppression activity matching that of healthy sleeping subjects (Fig. 1D and S2C). Notably, these sleep-like OFF-periods were never found when the same cortical areas were stimulated in awake healthy subjects (Fig. 1A and S2A). Why do awake brain-injured patients show cortical responses that are typical of the sleeping brain? A possibility is that structural lesions may lead to functional changes that precipitate intact portions of the thalamocortical system into pathological bistable dynamics *(40-42)*. For example, this might happen when subcortical lesions, such as diffuse axonal injury (DAI), interrupt a critical amount of fibers of the ascending activating systems *(43)*. In an extreme case, the thalamocortical system may be largely intact but functionally constrained to a pathological bistable state due a predominance of activity-dependent K+ currents *(35, 44, 45)*. Multifocal white matter lesions and DAI, may also induce bistability by engendering a state of cortico-cortical disfacilitation, that is by reducing recurrent excitation *(37)*. Indeed, intracellular recordings have shown that, following a surgical white matter undercut (cortical slab), pyramidal neurons can switch their discharge patterns from tonic firing to an intrinsically bursting regime promoting the alternation of up- and down-states *(41, 46–49)*.

In addition to altering intrinsic neuronal properties, a critical reduction of cortico-cortical excitation may shift the excitation/inhibition towards the latter. In physiological conditions, active inhibition (especially GABAB) plays an important role in inducing cortical OFF-periods *(18, 21)*. This is known to occur locally after a stroke *(50)*, but may involve the whole remaining cortex following severe, multifocal injury *(42)*.

Finally, multifocal brain injury can also induce critical functional shifts by altering the balance within the cortico-striatal mesocircuits *(51, 52)*. This mechanism is particularly relevant because it may lead to both cortical disfacilitation and thalamic hyperpolarization. Importantly, if the latter exceeds a given threshold, thalamic neurons may switch their firing pattern from tonic to bursting mode *(53)*, thus reinforcing bistable dynamics *(54)*.

Overall, different mechanisms, alone or in combination, may engender a tendency to bistable behaviours following brain injury. While their relative contribution is difficult to disentangle, it is worth noting that all the above mechanisms can be effectively engaged by a cortical perturbation. For example, a direct cortical hit with TMS may (i) trigger activity-dependent K+ currents and an OFF-period, if K+ channels are de-inactivated; (ii) massively recruit local inhibitory circuits leading to an OFF-period, if the excitation-inhibition balance is biased towards the latter; (iii) force hyperpolarized thalamo-cortical cells to fire bursts of action potentials back to the cortex and then fall into a prolonged silence, if thalamo-cortical cells are in a bursting mode. In fact, although the presence of slow waves in brain-injured patients has been shown by spectral analysis of the spontaneous EEG *(55–59)*, TMS perturbations could reveal the presence of OFF-periods in all patients, regardless of their prevalent background EEG pattern (Table S1) and even when slow waves were not evident in pre-stimulus ongoing activity (Fig. 2B, C and D).

## Pathological bistability, loss of complexity and loss of consciousness

What are the consequences of cortical bistability in brain-injured patients? An intriguing possibility is that this simple dynamic may impair both the differentiation and the integration of thalamocortical circuits, two properties that are thought to be jointly critical in sustaining the brain capacity for consciousness *(60, 61)*. In UWS patients OFF-periods were ubiquitously observed for TMS applied over parietal and frontal cortices (Fig. 1, Fig. 2 and Fig. S2). In this way, bistability obliterated the physiological differentiation of the impulse response in different cortical areas (i.e. the natural frequency; *(62))* normally observed in awake, conscious subjects. At the same time, OFF-periods curtailed deterministic interactions, as revealed by the correlation between the timing of their occurrence (SHFt) and the abrupt termination of phase-locked oscillations (PLFt). To the extent that recurrent interactions rely on the amplification of coherent activity across distributed sets of neurons, the scrambling of phases operated by the OFF-periods at each node may critically impair the emergence of large-scale cortical integration *(63)*.

In view of the above, here we assessed the relationships between the occurrence of OFF-periods, the duration of phase-locking and the temporal evolution of PCI, an index that is explicitly designed to quantify the joint presence of differentiation and integration in cortical networks *(12)*. In UWS patients the build-up of complexity (max PCIt) was shorter and never reached the levels attained in awake, conscious subjects; crucially, the time at which PCI stopped growing correlated significantly with both the occurrence of the OFF-period (maxSHFt) and the termination of phase locked-activity (PLFt)(Fig. 3). These results corroborate the hypothesis that bistability may be in a key position to impair overall brain complexity. Most important, these significant correlations draw a first link between neuronal events and global brain dynamics relevant for pathological loss and recovery of consciousness.

Along this line, a recent microscale study employing electrical stimulation and recordings in isolated cortical slices showed that phase-locking and complex causal interactions, as assessed by an adapted version of PCI, could be effectively restored by pharmacological interventions that reduce bistability and increase cortico-cortical excitability *(64)*. This microscale finding further suggests a causal link between bistability and complexity and may have translational implications since brain slices can be considered a simplified model of the electrophysiological state of cortical circuits under conditions of severe deafferentation.

## Clinical implications: recovering from pathological bistability

Loss of consciousness in UWS patients is associated with a variable degree of brain damage and physical disconnection of neural linkages *(65, 66)*. In a minority of brain-injured patients plastic structural changes, including axonal regrowth, may directly support behavioural recovery *(67, 68);* in others cases, functional adjustments may play a major role, while, the amount of structural brain damage remains substantially equal *(69-71)*. In this respect, to the extent that pathological sleep-like bistability represents a common functional endpoint disrupting large-scale interactions across structurally intact portions of the cortex, its reversal may potentially be relevant for clinical recovery.

The course of events illustrated in Fig. 5 is compatible with this hypothesis. This figure illustrates the results of longitudinal TMS/EEG measurements performed in one patient evolving from UWS to minimally conscious state (MCS), and eventually regaining consciousness. In this patient, behavioral recovery occurred in the space of two weeks and was associated with a progressive decrease of bistability and a concurrent recovery of causality and complexity.

**Fig. 5.**
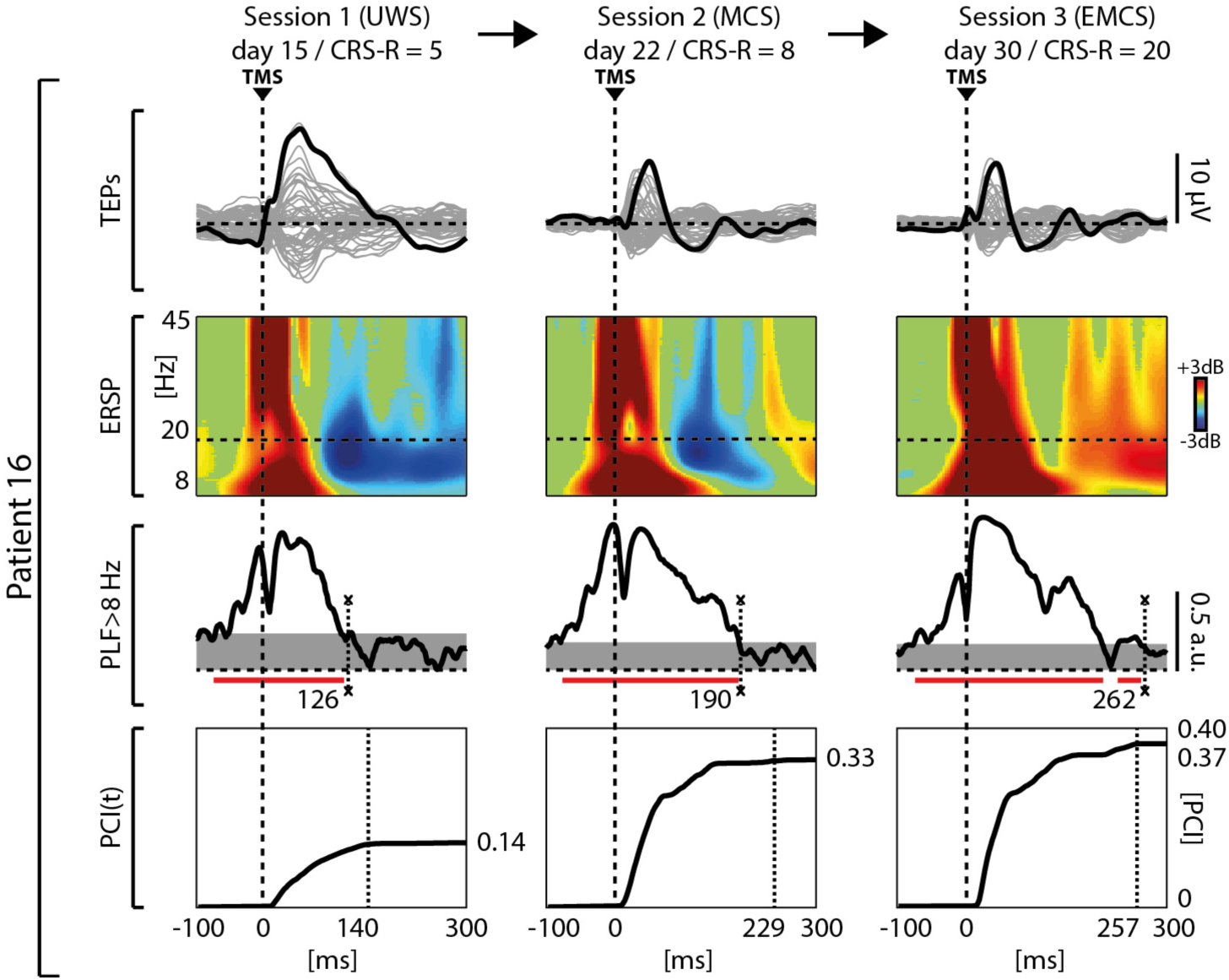
Longitudinal TMS/EEG measurements performed in one patient who gradually evolved from an unresponsive wakefulness syndrome (UWS), through a minimally conscious state (MCS) to emergence from a MCS (EMCS). Patient 16 (see Table S1) evolved from UWS to MCS, and EMCS; the first behavioral and TMS/EEG assessments (Session 1) were carried out 48 h after withdrawal of sedation, as patient exited from coma (*28*). The butterfly plot of the TMS-evoked potentials recorded from all 60 channels (grey traces), the corresponding ERSP and the PLF time course of the channel with the larger TEP are shown for each clinical status (UWS, MCS and EMCS) together with the temporal evolution of PCI. ERSP are color coded as in Fig. 1. The dashed horizontal line marks the 20 Hz frequency bin. The last significant (α < 0.01) time point in the PLF (above 8 Hz) is marked by a light-dashed vertical line. Time points above statistical threshold (grey shaded area) are underlined by a red horizontal line. The dashed vertical lines indicate the occurrence of the TMS pulse (bold dashed) and the time when PCI reaches its maximum value (max PCIt - light-dashed).

While elucidating the mechanisms of this recovery is clearly beyond of the scope of this work, the present observations raise important questions. What is the critical mass of neurons needed to recover from bistability in order to bring about detectable clinical changes? Does it matters whether this functional recovery occurs in a specific region of the cortex *(72)?* Can neuromodulation or pharmacological manipulation push neurons beyond the threshold for bistable dynamics thus allowing recovery of complexity? Are some patients just below this critical threshold? Different interventions *(73)*, such as zolpidem or amantadine administration *(74, 75)*, thalamic stimulation with Deep Brain Stimulation *(69, 76)* or low-intensity focused ultrasound pulsation *(70)*, transcranial direct current stimulation *(77, 78)* and vagal nerve stimulation *(79)* have demonstrated significant behavioural improvements in individual patients but a reliable read-out and interpretation of their end-point effects at the level of cortical circuits is still lacking. To the extent that sleep-like bistability represents the common functional endpoint of loss of complexity in severely brain-injured brains, detecting its presence and tracking its evolution over time, may offer a valuable read-out to devise, guide and titrate therapeutic strategies aimed at restoring consciousness. In this respect, it will be crucial to further elucidate the relationships between neuronal bistability and overall network complexity through extensive experiments across scales, species and models, spanning from ionic channel modeling to whole-brain simulations and macroscale measurements at the patient's bedside.

## Material and methods

### Study participants

Fifteen severely brain-injured patients took part in the present study (Table S1) by undergoing multiple behavioral assessments with the Coma Recovery Scale-revised (CRS-r; *(80)*) for a period of one week (four times, every other day), and one data recording session (TMS/EEG and rest EEG) within the same evaluation week. One additional patient (Patient 16 in Table S1) was clinically monitored for a period of 1 month and underwent three neurophysiological assessments: first in a UWS condition, then after evolution to a minimally conscious state (MCS), and eventually upon emergence from the minimally conscious state (EMCS). All UWS patients here reported were included in the low-complexity UWS subgroup described in a recent study *(10)*. As a control group, we enrolled twenty healthy volunteers (14 females; age range: 19-80 years), who underwent a general medical and neurological examination, in order to prevent potential adverse effects of TMS, and exclude major medical and/or neurological diseases as well as substance abuse.

### Experimental procedures

A single recording session was performed in each patient, except for the longitudinal assessment performed in Patient 16. During the recordings, UWS patients were lying in their beds with eyes open, and vigilance was continuously monitored. In case signs of drowsiness appeared, recordings were temporarily interrupted and patients were stimulated using the CRS-R arousal facilitation protocols *(80)*

All healthy subjects were recorded during wakefulness with eyes open while lying on a reclining chair with a headrest to ensure a stable head position. Eight healthy subjects also underwent an extra TMS/EEG measurement during NREM sleep (consolidated stage N3; *(81)*).

TMS targets were selected based on the individual anatomical MRI and the precision and reproducibility of stimulation were controlled using a Navigated Brain Stimulation (NBS) system (Nexstim Ltd., Finland).

As described in *(10)*, TMS targets were selected bilaterally within the frontal and the parietal cortices (Brodmann area - BA6 and BA7, respectively). The need to avoid direct stimulation of cortical lesions guided the specific selection of TMS targets *(30)* in UWS patients. Depending on the spatial extent and location of lesions in each individual patient, in the present study we considered the TMS/EEG measurements obtained by stimulating either one cortical site (BA6 or BA7) or both (see Table S1).

In healthy subjects, both BA6 and BA7 were targeted with TMS during wakefulness, while during NREM sleep only BA7 was targeted. This choice was dictated by the previous observation that during NREM sleep TMS evokes larger EEG responses in parietal areas as compared to frontal sites *(82)*.

Stimulation pulses were delivered with a Focal Bipulse figure-of-eight coil (mean/outer winding diameter ~50/70 mm, biphasic pulse shape, pulse length ~280 μs, focal area of the stimulation 0.68 cm^2^) driven by a Mobile Stimulator Unit (eXimia TMS Stimulator, Nexstim Ltd., Finland). Stimulation intensity was comparable between UWS patients (137.8±7.4 V/m) and healthy subjects (133.4±3.7 V/m; Wilcoxon ranksum test, p=0.62). For all the TMS/EEG measurements, the location of the maximum electric field induced by TMS on the cortical surface (hotspot) was always kept on the convexity of the targeted cortical gyrus with the induced current perpendicular to its main axis. In each TMS/EEG measurement, at least 200 stimulation pulses were delivered with an inter-stimulus interval randomly jittering between 2000 and 2300 ms (0.4–0.5 Hz).

All the experimental procedures were approved by the following ethical committees: Istituto di Ricovero e Cura a Carattere Scientifico Fondazione Don Gnocchi Onlus, Milan, Italy; Comitato Etico Interaziendale Milano Area A, Milan, Italy; Medicine Faculty of the University of Liège, Liège, Belgium. All healthy participants gave written informed consent, while for non-communicating UWS patients the informed consent was obtained by a legal surrogate.

### EEG recordings

EEG data were recorded using a TMS-compatible 60-channel amplifier (Nexstim Ltd, Finland), which gates the magnetic pulse artefact and provides artifact-free data from 8 ms after stimulation *(83)*. Raw recordings were referenced to a forehead electrode, online filtered between 0.1-350 Hz, and sampled at 1450 Hz. Two additional sensors were applied to record the electrooculogram (EOG). As previously recommended *(84)*, during all TMS/EEG recordings a masking sound was played via earphones and a thin layer of foam was placed between the coil and the scalp in order to abolish the auditory potentials evoked by the TMS loud click.

### Data analysis

Data analysis was performed using Matlab R2012a (The MathWorks Inc.). TMS/EEG recordings were visually inspected to reject trials and channels containing noise or muscle activity *(10, 30, 85)*. Then, EEG data were bandpass filtered (1-45 Hz, Butterworth, 3rd order), down-sampled to 725 Hz and segmented in a time window of ± 600 ms around the stimulus. Bad channels were interpolated using the spherical function of EEGLAB *(86)*. Recording sessions with either more than 10 bad channels or less than 100 artifact-free trials were excluded from further analysis. Then, trials were re-referenced to the average reference and baseline corrected. Finally, Independent Component Analysis (ICA) was applied in order to remove residual eye blinks/movements, TMS-evoked and spontaneous scalp muscle activations.

In UWS patients (n=16), we selected 24 TMS/EEG measurements (Table S1) for analyses (10 BA6 stimulation and 14 BA7 stimulation). In healthy awake subjects we performed 40 TMS/EEG measurements (20 BA6 stimulation and 20 BA7 stimulation). In addition, 8 TMS/EEG measurements were performed in a subset of healthy subjects during N3 stage of NREM sleep (BA7 stimulation).

Rest EEG recordings collected in UWS patients were evaluated according to a clinical classification recently proposed *(87)* after bandpass filtering between 1-70 Hz, downsampling to 725 Hz and re-referencing to the standard longitudinal montage. The EEG category of each patient is reported in Table S1. In the text, data are shown as means ± SEM. Group analyses were performed in MATLAB by non-parametric ANOVA with Kruskal-Wallis test and by Wilcoxon ranksum test.

### Amplitude of TMS-evoked slow wave and high-frequency suppression

We detected the occurrence of a cortical OFF-period by quantifying (1) the amplitude of a TMS-evoked slow wave (< 4 Hz) and (2) the significant suppression of high-frequency (> 20 Hz) EEG power compared to pre-stimulus *(13, 23-26)*. Operationally, for each EEG channel i (1 to 60), we followed the stepwise procedure presented in Fig. S1 and described below:

1. To assess the amplitude of TMS-evoked slow waves, single trials were low-pass filtered below 4 Hz (third order Chebyshev filtering as in *(13)*), re-referenced to the mathematically-linked mastoids, averaged and eventually rectified. For each channel *i*, the maximum Slow Wave amplitude (max SWa(i)) was computed as the maximum amplitude of the rectified signal within the 8-350 ms time window (Fig. S1A).
2. To assess the suppression of high-frequency (> 20 Hz) EEG power, we applied the event related spectral perturbation (ERSP) routine implemented in EEGLAB *(86)*. Specifically, single trials were time-frequency decomposed between 8 and 45 Hz using Wavelet transform (Morlet, 3.5 cycles; as in *(62)*) and then normalized with the full-epoch length (here ranging from −350 to 350 ms) single-trial correction *(85, 88)*. The resulting ERSPs were averaged across trials and baseline corrected (from -350 to -100 ms; *(88)*). Furthermore, power values that were not significantly different from the baseline (bootstrap statistics, α<0.05, number of permutations = 500) were set to zero. Then, the time course of the significant high-frequency EEG power was obtained by averaging over frequency the ERSP values above 20 Hz *(23)*.

Then, from the time course of significant high-frequency EEG power of each channel i, we extracted three morphological parameters: the integral between 100 and 350 ms of the high frequency (> 20 Hz) power (HFp(i)), the maximum value of high-frequency power suppression (max SHFp(i)) and the timing of the maximum high frequency (> 20 Hz) power suppression (max SHFt(i)).

All the measures described above (max SWa(i), HFp(i), max SHFp(i) max SHFt(i)) and calculated at the single channel level were averaged over the four channels closer to the stimulation site (Fig. S1D) as in *(89)* and the resulting averages were labelled: max SWa (Slow Wave amplitude), HFp (High Frequency power), max SHFp (maximum value of Suppression of High Frequency power), max SHFt (timing of the maximum value of Suppression of High Frequency).

### Phase-locking factor

The impact of the OFF-periods on local causal interactions was assessed by means of broadband (> 8 Hz) phase-locking factor (PLF; *(90, 91)*). Specifically, for each EEG channel *i* (1 to 60), single trials were high-pass filtered above 8 Hz (third order Butterworth filter) and PLF was computed as the absolute value of the average of the Hilbert Transform of all single trials. Assuming a Rayleigh distribution of the baseline values from -500 ms to -100 ms, PLF time points that were not significantly different from baseline (α<0.01) were set to zero. For each channel *i*, the latest significant PLF time point was identified and labelled as max PLFt(i). Finally, max PLFt (timing of the last significant time point of Phase Locking) was calculated as the average of max PLFt(i) over the four channels closer to the stimulation site (Fig. S1D).

### Perturbational Complexity Index

In order to assess the effects of bistable dynamics on the complexity of global causal interactions, we calculated PCI by applying the same automatic procedure described in two recent papers *(10,12)*. In summary, after source modeling, statistical analysis was performed to extract the significant spatiotemporal pattern of the TMS-evoked responses. Then, PCI was obtained as the Lempel–Ziv complexity of the matrix of significant cortical source activity and normalized by source entropy. To further study the relationships between bistable dynamics and the emergence of complex interactions, we used the temporal evolution of PCI, i.e PCI(t), which describes the buildup of complexity of the deterministic brain responses to TMS over time. Specifically, we rounded PCI(t) to the second decimal place and we measured the first time point in which PCI(t) reached its maximum (max PCIt).

